# Myelin lipids as nervous system energy reserves

**DOI:** 10.1101/2022.02.24.481621

**Authors:** Ebrahim Asadollahi, Andrea Trevisiol, Aiman S. Saab, Zoe J. Looser, Payam Dibaj, Kathrin Kusch, Torben Ruhwedel, Wiebke Möbius, Olaf Jahn, Myriam Baes, Bruno Weber, E. Dale Abel, Andrea Balabio, Brian Popko, Celia M. Kassmann, Hannelore Ehrenreich, Johannes Hirrlinger, Klaus-Armin Nave

**Author notes:** Correspondence Prof. Klaus-Armin Nave, PhD, Dept. of Neurogenetics, Max Planck Institute of Experimental Medicine, Hermann-Rein-Str. 3, 37075 Göttingen, Germany.

## Abstract

Neuronal functions and impulse propagation depend on the continuous supply of glucose^1,2^. Surprisingly, the mammalian brain has no obvious energy stores, except for astroglial glycogen granules^3^. Oligodendrocytes make myelin for rapid axonal impulse conduction^4^ and also support axons metabolically with lactate^5–7^. Here, we show that myelin itself, a lipid-rich membrane compartment, becomes a local energy reserve when glucose is lacking. In the mouse optic nerve, a model white matter tract, oligodendrocytes survive glucose deprivation far better than astrocytes, by utilizing myelin lipids which requires oxygen and fatty acid beta-oxidation. Importantly, fatty acid oxidation also contributes to axonal ATP and basic conductivity. This metabolic support by fatty acids is an oligodendrocyte function, involving mitochondria and myelin-associated peroxisomes, as shown with mice lacking Mfp2. To study reduced glucose availability *in vivo* without physically starving mice, we deleted the Slc2a1 gene from mature oligodendrocytes. This caused a significant decline of the glucose transporter GLUT1 from the myelin compartment leading to myelin sheath thinning. We suggest a model in which myelin turnover under low glucose conditions can transiently buffer axonal energy metabolism. This model may explain the gradual loss of myelin in a range of neurodegenerative diseases^8^ with underlying hypometabolism^9^.

## Main

Oligodendrocytes are glycolytic cells that can provide myelinated axons with lactate or pyruvate for the generation of ATP^5,6,10^. Surprisingly, Drosophila revealed that the metabolic support of axons by associated glia preceded the evolution of myelin in vertebrates^11,12^. In non-myelinating species, notably in Drosophila larvae^13,14^ and in lamprey^15^, the axon-associated glial cells that lack myelin accumulate lipid droplets, which are well-known energy reserves^16,17^. We therefore hypothesized that lipid droplets, which are not a feature of mammalian oligodendrocytes^18^, were ‘replaced’ in vertebrate evolution by the exuberant synthesis of lipid-rich membranes as myelin^19^. Spirally wrapped around axons, these myelin membranes enable saltatory impulse propagation^20^ and block ephaptic coupling between neighboring axons^12^. If myelin had indeed replaced lipid droplets, we reasoned it is possible that myelin has retained a function in energy metabolism.

### Glial cell survival in the absence of glucose

Cultured cells can die rapidly without glucose^21,22^. We therefore asked whether myelinating oligodendrocytes, when glucose deprived, can metabolize myelin lipids for survival. We chose the acutely isolated optic nerve (Fig.1a) as a model system, in which glial cells survive when provided with glucose (or lactate) and oxygen. We analyzed fully myelinated transgenic mice at age 2 months, expressing fluorescent proteins in mature oligodendrocytes (*Cnp-mEos2-PTS1*)^23^ or astrocytes (*Aldh1L1-GFP*). Optic nerves were incubated at 37°C in artificial cerebrospinal fluid (aCSF) containing either 10 mM glucose, 0 mM glucose (see methods section) termed ‘glucose-free’, or in low glucose (termed ‘starved’). After 24 hours, the total number of cells (DAPI positive), the number of apoptotic cells (propidium iodide positive), and the identity of surviving cells were determined by fluorescence analysis of sectioned nerves. Surprisingly, the large majority (>97%) of oligodendrocytes appeared healthy after 24h in glucose-free medium, whereas more than 70% of astrocytes had died (Fig.1b-f). Oligodendrocyte precursor cells (OPC) and microglia were also not reduced. Next, we compared earlier time points and found no cell death at 16h (Fig.1c). This suggests that all glial cells in the myelinated optic nerve can survive in the absence of glucose by utilizing a preexisting energy reserve.

**Fig 1.**
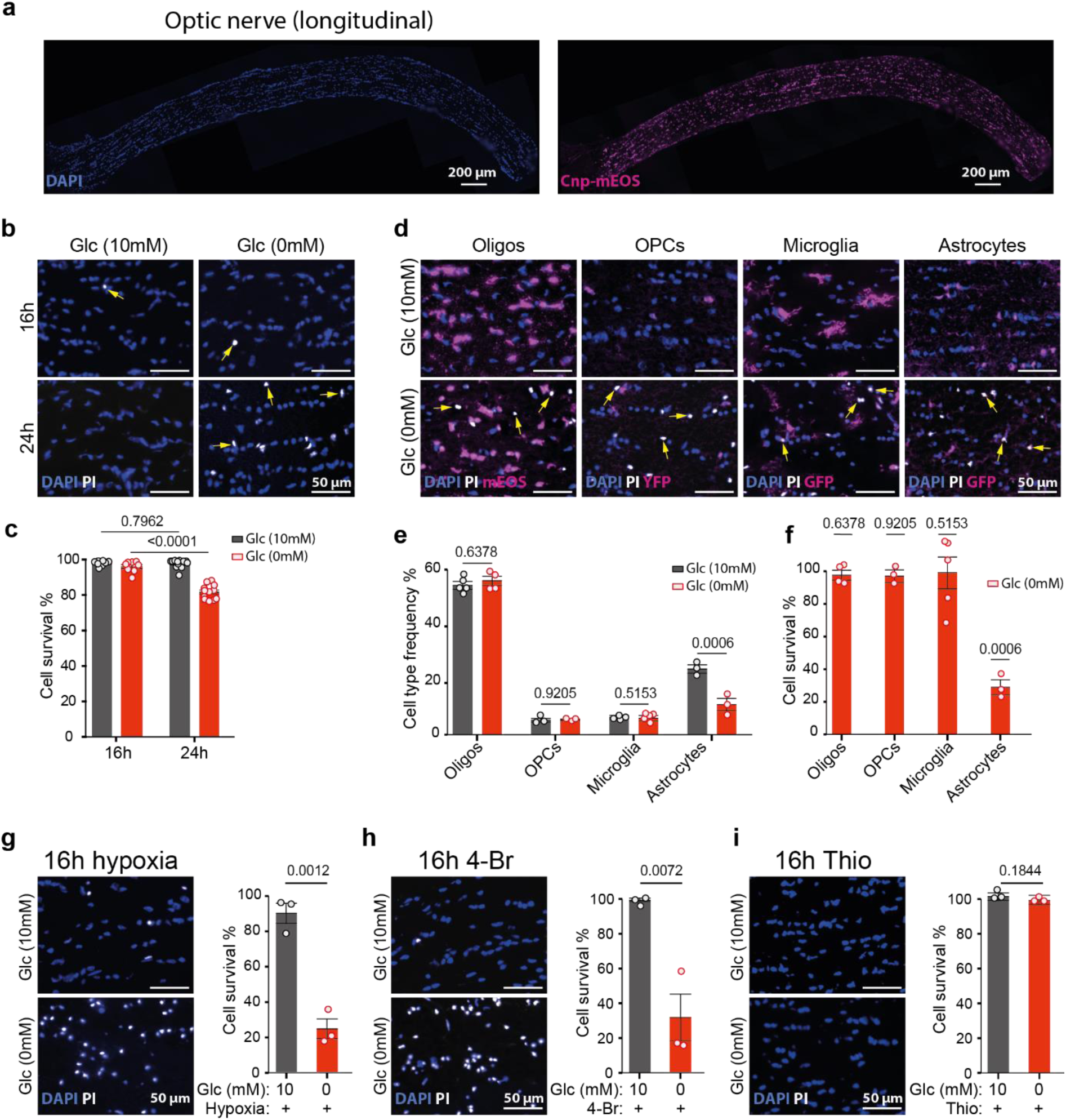
Survival of glial cells in the absence of glucose requires the utilization of fatty acids. **a**, Myelinated optic nerve from a *Cnp-mEOS* reporter mouse (age 2 months), acutely isolated and maintained *ex vivo*. **Left:** longitudinal section showing all cell nuclei labeled by DAPI (blue). **Right:** oligodendrocytes labelled by the expression of mEOS (magenta). **b**, Higher magnification of optic nerve glia (DAPI: blue), with dying cells (yellow arrows) taking up propidium iodide (PI: white). **Left:** virtually all cells survive in the presence of 10 mM glucose (Glc). **Right:** in the absence of glucose (0 mM glucose/10 mM sucrose), cells survive up to 16 hours but many die at 24h, as quantified in **c**. **c**, Cell survival in optic nerves incubated in the presence of glucose (grey) or in the absence of glucose (red) for 16h and 24 h, quantified by subtracting the dying cells (PI+DAPI+) from total number of cells (DAPI+) (n=9-12; average ±SEM, p-values as indicated; data from d-f were included). **d**, Different vulnerabilities of the major glial subtypes to 24h glucose withdrawal. Optic nerve longitudinal sections were colabeled by DAPI (quantifying cell number), PI (dying cells), and genetically expressed fluorescent markers for oligodendrocytes (Oligos: Cnp-PTS1-mEOS), oligodendrocyte precursor cells (OPC: Ng2-YFP), microglia (Cxcr1-GFP) and astrocytes (Aldh1l1-GFP). Note that in the absence of glucose, cell nuclei are shrunken in size and oligodendrocytes (magenta, far left panel) show no overlap with dying cells (white), as quantified below. **e**, Frequency of each cell type in the optic nerve after 24h incubation in the presence (10 mM) or absence (0 mM) of glucose (same data as in d; n=3-5; average ±SEM, p-values as indicated). **f**, Same data as in (e), survival rate of glial cell subtypes in the optic nerve after 24h incubation in the absence of glucose (0 mM), normalized to the respective survival in glucose containing aCSF (100%; n=3-5; average ±SEM, p-values as indicated). **g**, Under an N2 atmosphere, many cells in the optic nerve cope by anaerobic glycolysis and survive for 16h if glucose is present (top), whereas glucose-deprived cells largely die (bottom), demonstrating the need for oxidative phosphorylation to utilize the endogenous energy reservoirs. **h**, Application of 4-Bromo-crotonic acid (4-Br), an inhibitor of mitochondrial beta-oxidation, causes widespread cell death (PI-positive) if the optic nerve is glucose deprived (0 mM), but not in controls (10 mM). This demonstrates that the critical energy reserves of the optic nerve are derived from fatty acids, and that 4-Br is not cytotoxic by itself. **i**, Application of Thioridazine (Thio), an inhibitor of peroxisomal beta-oxidation, does not cause glial cell death in glucose-deprived nerves, suggesting that for glial cell survival mitochondrial beta-oxidation is sufficient. All animals were at two months of age and percentages (in g-i) were calculated relative to overall survival after 16h with glucose under normoxia (in c). Error bars: mean+/-SEM (t-test).

In the presence of 1 mM glucose, a concentration insufficient to maintain axonal conduction (Suppl Fig.3a), all cells of the optic nerve stayed alive for at least 24h. Glucose is essential for the pentose-phosphate pathway and the synthesis of nucleotides. To rule out that glucose-free medium is detrimental independent of the reduced energy metabolism, we incubated optic nerves in aCSF containing as little as 1.5 mM beta-hydroxy butyrate as an alternate energy source and detected no cell death after 24h (Suppl Fig.1a, b). Moreover, optic nerves that were kept glucose-free in the presence of a ROS inhibitor (S3QEL-2) and a mitochondrial ROS scavenger (MitoTEMPO) did not show enhanced cell survival, suggesting that cell death is not caused by the generation of ROS (suppl Fig.1c, d).

Our hypothesis that (myelin) lipids provide the energy reserve for respiratory ATP generation was supported by experiments, in which optic nerves were incubated for only 16 hours in glucose-free aCSF in combination with severe hypoxia (N2 atmosphere). Here, cell death was extensive with 76% propidium iodide labelled cells, suggesting that all cell types are affected (Fig.1g and data not shown). Thus, in the absence of glucose virtually all glial cells appear to have survived by oxidative phosphorylation.

### Myelin-derived fatty acids as energy reserves

We next asked whether lipids comprise the postulated energy reserve for oxygen-dependent generation of ATP via the beta-oxidation of fatty acids. Optic nerves were incubated without glucose and under normoxia, but in the presence of 25 µM 4-bromo-crotonic acid (4-Br), a thiolase inhibitor of mitochondrial fatty acid beta oxidation and ketolysis. Application of this drug dramatically reduced cell survival to 30% at 16h (Fig.1h). Importantly, 4-Br had no effect on glial survival in the presence of glucose (Fig.1h), ruling out unspecific toxicity. Next, we tested 5 µM Thioridazine (Thio), an inhibitor of peroxisomal beta-oxidation^24^ and also claimed to block mitochondrial beta-oxidation^25^. However, Thio had no obvious effect in the absence of glucose (Fig.1i). This suggests that its effect on peroxisomes is distinct or at least partially compensated by mitochondrial beta-oxidation (but see below).

If myelin-derived fatty acids were the main source of metabolic energy, starvation might lead to a visible loss of myelin membranes. To directly determine this, we maintained optic nerves in low glucose (0.5 mM) or regular (10 mM glucose) medium and analyzed them 24 hours later (i.e. before major glial cell death; Fig.1c and data not shown) by electron microscopy and quantitative morphometry (Fig.2a-c). Starvation led to increased g-ratios (Fig.2a-c). However, these observations were not conclusive because starvation also caused unspecific swellings of the axonal and myelin compartments. Compatible with active demyelination was the emergence of vesicular structures underneath the myelin sheaths in starved nerves (Suppl Fig.2a, b), likely caused by autophagy (see further below).

**Fig 2.**
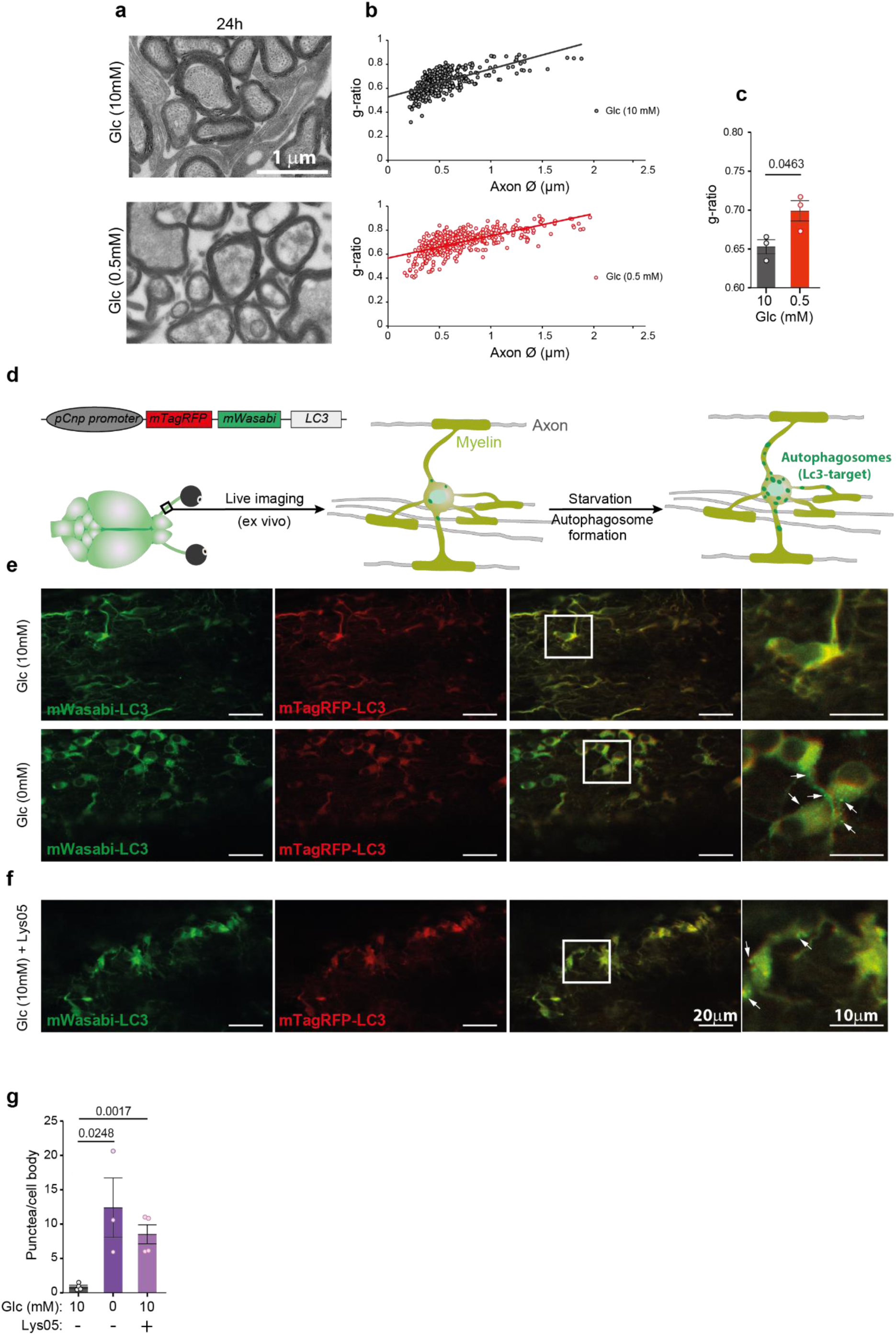
Glucose-deprivation of the optic nerve causes shrinkage of the myelin compartment. **a**, Electromicroscopic images of optic nerve cross-sections (wild type mice; age two months), taken after 24h of incubation in medium with 10 mM glucose (top) or 0.5 mM glucose (bottom). **b**, Scatter plot of calculated g-ratios (outer fiber diameter/axon diameter) as a function of axon caliber, revealing myelin loss when optic nerves are exposed to low glucose (red dots, bottom) in comparison to 10 mM glucose containing medium (black dots, top). **c**, Bar graph with calculated mean g-ratios (same data as in b). (n=3 for both 0 mM and 10 mM; Error bars: mean+/-SEM; t-test). **d**, Schematic depiction of the *pCNP-mTagRFP-mWasabi-LC3* transgene used for oligodendrocyte-specific expression of LC3 fusion protein. The two fluoropores (red and green) give LC3 a yellow color and reveal diffuse cellular distribution. Note that mTagRFP-mWasabi-LC3 becomes an autophagosome-specific marker, when LC3 is translocated to the membrane of newly formed autophagophore/autophagosomes (green puncta). **e**, Live imaging of optic nerves (*ex vivo*) from a *mTagRFP-mWasabi-LC3* transgenic mouse, incubated in aCSF with 10 mM (top) or 0 mM (bottom) glucose. Note the ubiquitous expression of the LC3 fusion protein. Specific labeling of autophagosomes (green puncta, arrows on right image) occurs only in the absence of glucose with an all-or-none difference. **f**, Control experiment, revealing the accumulation of autophagosomes also in oligodendrocytes in the presence of 10 mM glucose after applying the specific inhibitor, Lys05 (10 µM). **g**, Quantification of the data in e-f, normalized to the number of cell bodies (Adult mice, 2-5 months old; n=4 for 10 mM glc+/-Lys05 and n=3 for 0 mM glc; error bars: mean +/-SEM; t-test).

We also subjected entire optic nerves to quantitative proteome analysis following either 16h in glucose-free medium (when all cells survive), or 24h in 1 mM glucose-containing medium, both in comparison to a regular medium (10 mM glucose). Interestingly, we noticed a non-significant trend towards higher myelin protein abundance (Suppl Fig.2c-e), most likely because autophagy liberates proteins from compact myelin that are otherwise not solubilized. In these nerve lysates, the abundance of some glycolytic enzymes was reduced (e.g. PFKAM) whereas fatty acid binding proteins (FABP3) and enzymes of fatty acid metabolism (ACAD9) were increased. A higher steady-state level of some autophagy-related proteins was only detected when nerves were maintained in low (1 mM) glucose (Suppl Fig.2e), most likely because low glucose is required for RNA synthesis. Importantly, after 16h in glucose-free medium also western blotting revealed a significant increase of ACAT1 and BDH1, enzymes involved in fatty acid and ketonbody metabolism, respectively (Suppl Fig.2f, g).

### Glucose withdrawal increases markers of autophagy

In adult mice, food withdrawal induces LC3 positive autophagosomes in neuronal perikarya but not in axons^26^. Is myelin degradation nevertheless mediated by autophagy? To study autophagy in optic nerves, we generated a new line of *pCNP-mTagRFP-mWasabi-LC3* transgenic mice that express a tandem (pH-sensitive) fluorescent tag^27^ in oligodendrocytes (Fig.2d). Indeed, 8.5 hours after glucose withdrawal from optic nerves we observed the accumulation of autophagosomes in oligodendrocytes (Fig.2e, g). In glucose containing medium (10 mM), however, these organelles were only detectable when their degradation was specifically inhibited, e.g. by Lys05 (Fig.2f, g). This suggests that in oligodendrocytes a basal level of autophagy already exists that strongly increases upon glucose deprivation. This may explain why transcriptional upregulation of autophagy genes by its master regulator TFEB is not required for utilizing fatty acids from the myelin compartment (see further below), as evidenced by the unaltered cell survival of glucose-deprived optic nerves from oligodendrocyte-specific TFEB cKO mice (Suppl Fig.2h, I).

### Lipid metabolization supports axon function

Since axonal conduction is energy-dependent, we asked next whether myelin-derived fatty acids can also support the axonal compartment under low glucose conditions. As a readout for axon function in acutely isolated optic nerves, we determined the size of the evoked compound action potential (CAP) after electrical stimulation^7^ (Fig.3a, b). These recordings were performed in combination with real-time monitoring of the axonal ATP levels in the same nerves, using a genetically encoded ATP sensor expressed in the axonal compartment^10^ (Fig.3c).

**Fig 3.**
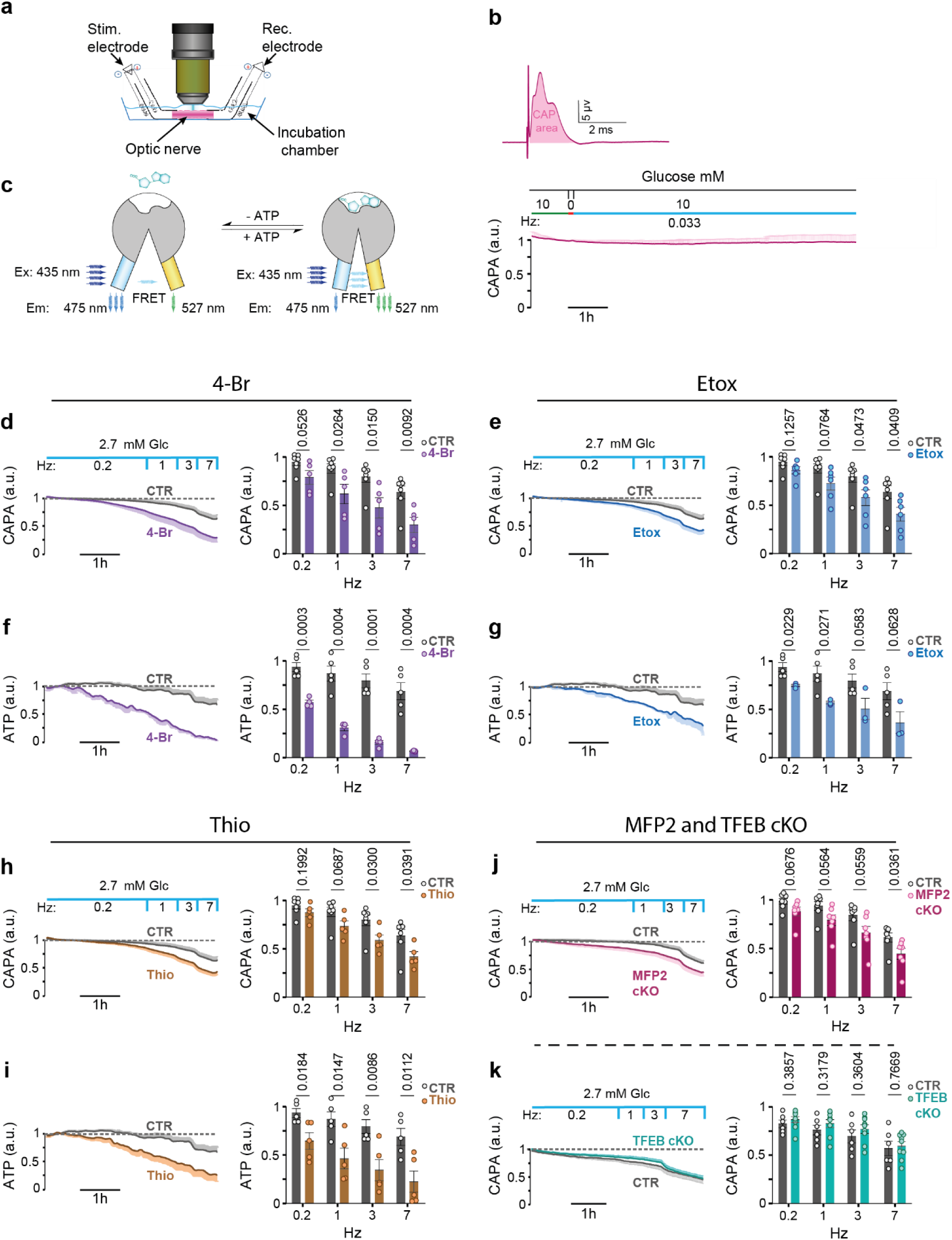
Fatty acid beta-oxidation in oligodendrocytes supports axonal energy metabolism and function. **a**, Schematic drawing of the *ex vivo* set-up used for stimulating and recording optic nerve compound action potentials (CAP) and the simultaneously microscopic monitoring of axonal ATP levels by ratiometric FRET analysis. **b, top:** CAP recording from an optic nerve maintained in 10 mM glucose with the ‘CAP area’ (CAPA) shaded in red below the curve. **bottom:** Stable CAPA, normalized to 1.0, in the presence of 10 mM glucose and normoxia during low spiking frequency (1 per 30s). Note the 5min glucose withdrawal (‘0’) to partially deplete astroglial glycogen stores. **c**, Schematic depiction of ratiometric FRET analysis, using the ATP sensor ATeam1.03^YEMK^ expressed in the neuronal/axonal compartment of transgenic mice^10^. **d**, Optic nerves, maintained functionally stable with a reduced glucose concentration (2.7 mM) and minimal spiking activity (0.2 Hz), were exposed to 4-Bromo-Crotonic acid (4-Br, 25 µM, n=5), an inhibitor of mitochondrial fatty acid beta-oxidation. Note the progressive decline of optic nerve conductivity beginning at 0.2 Hz, quantified by comparing the CAPA at increasing spiking frequencies to low glucose controls (CTR, n=7). **e**, Optic nerves, maintained at 2.7 mM glucose with increasing stimulation frequency, were exposed to Etomoxir (Etox, 5 µM, n=6), an inhibitor of long chain fatty acid uptake into mitochondria. Note the faster declining CAPA in comparison to controls (n=7). **f**, Same as in (d), demonstrating a progressive loss of the steady-state axonal ATP recorded by FRET analysis. Note the faster and stronger effect of inhibiting fatty acid beta-oxidation on the axonal ATP level at 0.2Hz stimulation (n=4) compared to controls (n=5). **g**, Analysis of axonal ATP in Etox-treated nerves (n=3) and controls (n=5) as before. **h**, Optic nerves, maintained at 2.7 mM glucose with increasing stimulation frequency, were exposed to Thioridazin (Thio, 5 µM, n=5), an inhibitor of peroxisomal fatty acid beta-oxidation. Analysis of faster declining CAPA (in h) in comparison to controls (n=7). Note the difference to cell survival (Fig.1i), which does not depend on peroxisomal functions. **i**, Analysis of axonal ATP in Thio-treated nerves (n=5) and controls (n=5) as before. **j**, Optic nerves from *Cnp-Cre*^*+/-*^*::Mfp2*^*flox/flox*^ mice, lacking peroxisomal beta-oxidation in oligodendrocytes^29^, and control mice, maintained at 2.7 mM glucose with increasing stimulation frequency (n=7 each). The stronger CAPA decline in mutant nerves (7Hz) confirms the oligodendrocyte-specific contribution to axonal conduction blocks. **k**, Optic nerves from *Cnp-Cre*^*+/-*^*::Tfeb*^*flox/flox*^ mice, lacking a key transcription factor for autophagy from oligodendrocytes, and control mice, maintained at 2.7 mM glucose with increasing stimulation frequency. Note that mutant nerves (n=9) are indistinguishable from controls (n=7) at any frequency, suggesting that fatty acid mobilization in oligodendrocytes does not depend on *de novo* autophagy induction but rather on ongoing myelin lipid turnover. All nerves were from mice aged 2 months. Bar graphs are mean values ±SEM (t-test) of data recorded during the last 5 min in each stimulation frequency step (controls in d and f were used for CAP and ATP measurements in different experiments(d-I)).

We first determined empirically the (set up-specific) threshold level of glucose concentration (Suppl Fig.3a, b), at which acutely isolated optic nerves, following 5min glycogen depletion, remained sufficiently energized to maintain a low firing rate (0.2 Hz) for 2 hours without decline of CAP area (Fig.3d; here: 2.7 mM glucose in aCSF). A subsequent gradual increase of the stimulation frequency (to 1 Hz, 3 Hz, 7 Hz) caused a gradual decline of the CAP, i.e. an increasing fraction of axons with conduction blocks. Both, preservation and decline of axon function could be quantified by calculating the curve integral, with the CAP areas (CAPA) plotted as a function of time. Importantly, when recordings were done in the presence of 4-Bromo-Crotonic acid (4-Br), an inhibitor of thiolase (Fig.3d), or Etomoxir (Etox), an inhibitor of mitochondrial carnitine palmitoyltransferase (CPT1) (Fig.3e), these blockers of mitochondrial fatty acid beta-oxidation caused a much more rapid decay of CAPA, i.e. loss of conductivity. When applied in the presence of 10 mM glucose these drugs had no toxic effects (Suppl Fig.3c-f). Under all conditions, we also monitored axonal ATP levels, which revealed strong parallels to the electrophysiological recordings (Fig.3f, g). Thus, when glucose is limiting the functional integrity of spiking axons is supported by fatty acid degradation.

To confirm that the decline of CAP, as observed under starvation, was not due to ROS production, we repeated our recordings in the presence of ROS inhibitors and scavengers. Indeed, these drugs could not “rescue” the CAP decline (Suppl Fig.3i, j).

To also rule out the possibility that inhibition of beta-oxidation interferes with degradation of toxic fatty acids generated in hyperactivated neurons^28^ and consequently the decline of the CAP, we compared optic nerve conduction also in the presence of 10 mM glucose. Indeed, high frequency (5-20 Hz) conduction of these nerves remained the same in the presence or absence of 4-Br (Suppl Fig.3k, l).

Next, we applied 5 µM Thioridazine (Thio), an inhibitor of peroxisomal beta-oxidation. Interestingly (and different from cell survival assays; Fig.1i), in the electrophysiological experiments similar results were obtained for the mitochondrial inhibitor and the peroxisomal inhibitor (Fig.3h, i). This allowed us to genetically confirm also the specific role of oligodendrocytes in axonal energy metabolism, by studying *Cnp-Cre*^*+/-*^*::Mfp2*^*flox/flox*^ mutant mice, in which oligodendrocytes selectively lack peroxisomal fatty acid beta-oxidation^29^. Indeed, increasing the axonal spiking frequency in these nerves to 7 Hz caused similarly the enhanced loss of CAPA as seen in Thioridazine treated nerves (Fig.3j). This observation was made in the absence of any underlying structural abnormalities of *Cnp-Cre*^*+/-*^*::Mfp2*^*flox/flox*^ optic nerves, including the number of axons, the number of unmyelinated axons, the number of ultrastructural axonal defects, and the number of microglia. Also, basic electrophysiological properties of the nerves, including excitability and nerve conduction velocity, were unaltered (Suppl Fig.3m-t). This demonstrated directly the role of oligodendrocytes in the support of starving axons, with a significant role of beta-oxidation in peroxisomes, many of which reside in the myelin compartment^23^.

Is axonal conductivity in starved nerves rescued by the upregulation of autophagy in oligodendrocytes? TFEB activates autophagy-related genes and lysosomal functions^30^ also in oligodendrocyte lineage cells^31^. Thus, we generated *Cnp-Cre::TFEB*^*flox/flox*^ mice for oligodendrocyte-specific ablation, but found that mutant optic nerve conduction remains indistinguishable from wildtype nerves with respect to conduction velocity (not shown) and the *ex vivo* CAP decline (Fig.3k). In contrast, in wildtype nerves under limiting glucose concentrations application of the autophagy inhibitor Lys05 caused a faster CAP decline (Suppl Fig.4a, b), suggesting some contribution of preexisting autophagy to axonal energy metabolism. In turn, the autophagy inducer 3,4-Dimethoxychalcone (DMC), an activator of TFEB and TFE3, improved nerve function, but failed to do so in nerves from *Cnp-Cre::TFEB*^*flox/flox*^ mice (Suppl Fig.4c, d). Taken together, these observations suggest that autophagy exists in oligodendrocytes, and that the master regulator TFEB is not critical for the mobilization of fatty acids. We also detected upregulation of autophagy in brain lysates of mice with reduced glucose availability to oligodendrocytes *in vivo* (see below and Fig.4i, j).

**Fig 4.**
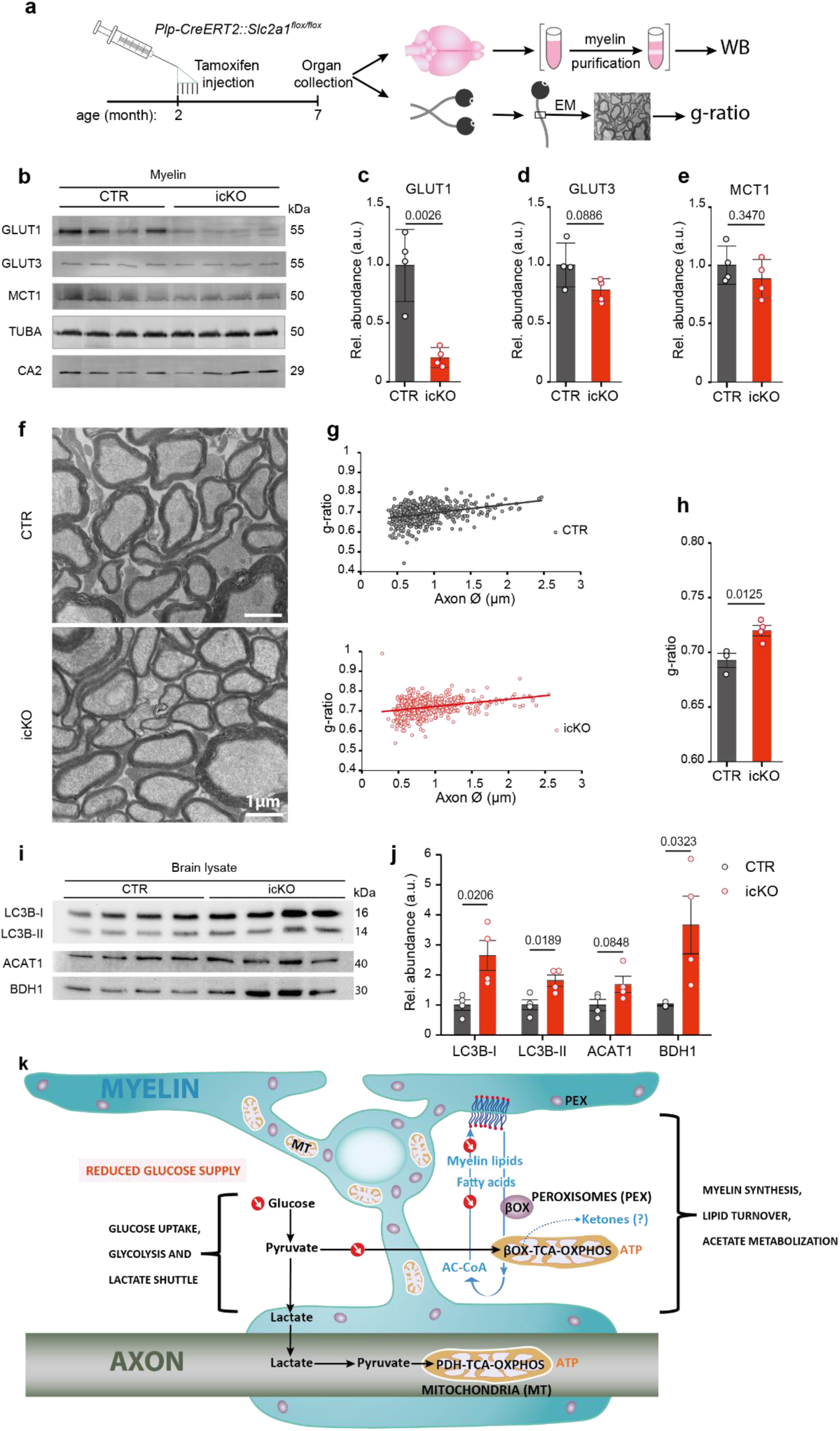
An *in vivo* model of reduced glucose availability causing myelin loss. **a**, To study the effect of reduced glucose availability on myelination *in vivo*, but without physically starving mice, we targeted the expression of glucose transporter GLUT1 selectively in oligodendrocytes. **Left:** schematic view of treating *Plp-CreERT2::Slc2a1*^*flox/flox*^ mice with tamoxifen at age 2 months for biochemical and ultrastructural analysis 5 months later. **Right:** decline of GLUT1 as quantified by Western blot (WB) analysis of purified myelin membranes; demyelination of the optic nerve quantified by electron microscopy (EM). **b**, Western blot analysis of myelin membranes purified from whole brain lysates five months after tamoxifen injection. Note the decrease of oligodendroglial GLUT1, but not neuronal GLUT3, or glial MCT1 (quantified in **c-e**). TUBA, α-tubulin; CA2, oligodendroglial carbonic anhydrase CA2. **f**, Electron micrographs of optic nerve cross section from GLUT1 conditional mutant (icKO) and control (CTR) mice, obtained five months after tamoxifen-induced recombination. Note the thinning of myelin but absence of axonal degeneration. **g**, Scatter plot of calculated g-ratios (fiber diameter/axon diameter) from optic nerve EM data, with regression lines as a function of axon diameter. **h**, By g-ratio analysis reduced glucose availability causes gradual demyelination in GLUT icKO mice (n=4) compared to controls (n=3). Error bars: mean+/-SEM (t-test). **i**, Western blots using brain lysate from GLUT1 icKO mice. **j**, Quantified blots (in i) normalized to total protein input as determined with fast green staining. (n=4 for control and n=4 for icKO, brains isolated 5 months after tamoxifen injection; t-test, mean +/-SEM). **k**, Proposed Working model of glycolytic oligendrocytes with a myelin compartment that constitutes a lipid-based energy buffer. During normal myelin turnover, the degradation of myelin lipids in lysosomes (Lys) liberates fatty acids (FA) for beta-oxidation (ß-Ox) in mitochondria (MT) and peroxisomes (PEX) leading to new myelin lipid synthesis. When glucose availability is reduced, as modeled in GLUT1 icKO mice, myelin synthesis drops and instead fatty acid-derived acetyl CoA begins supporting mitochondrial respiration for oligodendroglial survival. Such a shift of normal myelin turover to lipid-based ATP generation allows oligodendrocytes to share more (glucose-derived) pyruvate/lactate with the axonal compartment to support ATP generation and prevent axon degeneration. Note that glucose is never absent *in vivo* and that myelin-associated peroxisomes^23^ are better positioned than mitochondria to support the axonal compartment with the products of fatty acid beta-oxidation. Whether oligodendrocytes also use ketogenesis to metabolically support axons and other cells is not known.

### Oligodendroglial glucose deprivation *in vivo*

Acute glucose deprivation in short *ex vivo* experiments can provide *proof of principle* for the utilization of myelin lipid-derived energy. However, this system cannot model longer-lasting phases of hypoglycemia that can occur in real life, e.g. upon starvation. We therefore searched for *in vivo* evidence that oligodendroglial glucose deprivation, when modeling e.g. prolonged hypoglycemia, causes detectable loss of myelin. Real ‘starvation’ experiments are not possible and have difficult to control side effects, such as ketosis. To circumvent this, we generated a line of tamoxifen-inducible *Plp1*^*CreERT2/+*^*::Slc2a1*^*flox/flox*^ conditional mutant mice, which allowed us to eliminate GLUT1 expression specifically in mature oligodendrocytes, i.e. by tamoxifen administration at the age of two months (Fig.4a). We expected a slow decline of this glucose transporter, because GLUT1 is associated with myelin^7^ and should have the slow turnover of myelin structural proteins. Moreover, *Plp1*^*CreERT2*^ recombination efficacy is <100%. Thus, five months after tamoxifen administration we found a significant but still incomplete decrease of GLUT1 on Western blots of purified myelin. In contrast, GLUT3, MCT1, α-tubulin, and oligodendroglial carbonic anhydrase-2 were unaltered in abundance (Fig.4b-e).

Indeed, mutant animals showed no obvious behavioral defects and lacked visible neuropathological changes (Suppl Fig.5a-c). However, when the myelin sheath thickness was quantified by electron microscopy (Fig.4f), g-ratio analysis of the optic nerve revealed significant loss of myelin membranes in the absence of obvious axonal pathology (Fig.4 g, h; Suppl Fig.5d-i). At the same time, there were no signs of inflammation or altered electrophysiological properties (suppl Fig5.j-m). Western blot analysis of mutant brain lysates showed elevated levels of LC3b I, II, BDH1 with a tendency for more ACAT1 (Fig.4i, j). These *in vivo* observations strongly support the idea that myelin is gradually metabolized when oligodendrocytes lack normal glucose uptake, and that the mechanism of myelin loss is the catabolic arm of normal myelin turnover.

## Discussion

Our *in vitro* and *in vivo* data, when combined, lead to to a new working model of myelin, which not only enables fast axonal impulse propagation, but also provides a local energy reserve when circulating glucose is very low. This model (summarized in Fig.4k) thus extends the model of glycolytic oligodendrocytes providing pyruvate or lactate as metabolic support to spiking axons^5–7,10^.

In the present study, a key experiment was the direct analysis of both axonal conductivity and ATP levels in the myelinated optic nerve under defined metabolic conditions and in the presence of specific metabolic inhibitors, complemented by oligodendrocyte-specific gene targeting experiments *in vivo*. The latter allowed us to confirm a gradual loss of myelin membranes in response to reduced cellular glucose availability. We hypothesize that in the absence of continued myelin synthesis due to glucose deprivation, the ongoing myelin turnover and the constitutively active pathway of myelin catabolism become diverted to support mitochondrial energy metabolism. In fact, this might be the fastest utilization of readily available metabolic energy.

Under physiological conditions, cellular energy deprivation can be caused by transient hypoglycemia, but not by aglycemia (except for ischemic stroke). Thus, in starved white matter tracts myelin-derived fatty acids would only have to compensate for some decrease of glucose availability. In oligodendrocytes, fatty acid beta-oxidation in mitochondria creates acety-CoA and can directly support the TCA cycle and respiration, thereby sparing glycolytic pyruvate/lactate for axonal metabolic support. However, the transfer of energy-rich metabolites could be more complex. Oligodendrocytes are able to generate acetoacetate from acetyl-CoA and acetoacetyl-CoA in the cholesterol pathway (by HMG-CoA lyase or direct deacylation)^32,33^. This pathway fits well recent findings in Drosophila, in which glycolytically impaired glial cells switch to beta-oxidation to support neuronal metabolism with lipid-derived ketone bodies^17,34^. In mammalian brains, ketone bodies increase with age^35^ and can spread horizontally like lactate through monocarboxylate transporters^36^. Moreover, myelin-derived metabolites, including amino acids and free acetate, can pass via gap junctions from oligodendrocytes to all other glia in the ‘pan-glial’ syncytium^37^. This might explain why in glucose-deprived optic nerves also glial cells that lack a myelin compartment survive under aerobic condition. Mice with cell type-specific deletions of MCT1^38^, connexins and pannexins, will help define these pathways in the future. The higher vulnerability of astrocytes to glucose deprivation is thus puzzling and may reflect an irreversible metabolic switch to glycolysis (and thus glucose dependency). Our experiments with specific inhibitors suggest that possible toxic effects of ROS, as studied in astrocytes^39^, are less likely.

Regardless of the possible downstream mechanism, our principle finding that myelin lipids are glial energy reserves, metabolically coupled to the neuronal compartment, could change the view on ‘white matter’. Myelin lipids vastly exceed the amount of astroglial glycogen, which can support neurotransmission and spiking axons, but is (experimentally) depleted within minutes. Unlike glycogen breakdown, the beta-oxidation of myelin-derived fatty acids does not support rapid axonal spiking. Thus, severe hypoglycemia will cause conduction blocks, such as in patients treated with insulin^2^. However, the utilization of myelin-derived energy as a buffer may prevent further decline of ATP and irreversible axonal degeneration in the CNS^40^.

If nutritional stress becomes prolonged, myelin energy reserves will also reach a limit. We note reports of MRI-detectable white matter lesions in patients with diabetic hypoglycemic coma^41^ or severe anorexia nervosa^42,43^, which was mechanistically unexplained. Individuals with obesity that underwent gastric bypass (bariatric) surgery have developed encephalopathy and peripheral neuropathy^44^. Earlier reports of physically starved rats^45^ showed also peripheral demyelination and it is likely that Schwann cells that contain glycogen granules^46^ and oligodendrocytes have similar functions in axonal metabolic support^5,9,47^.

Our findings have also relevance for neurodegenerative diseases. In multiple sclerosis (MS), axon loss has been attributed to an energy failure^48^ which could further drive demyelination. Many neuronal disorders, including Alzheimer’s disease, have been associated with hypometabolism^9^ and white matter abnormalities^8,49,50^. Even psychiatric diseases have exhibit unexplained myelin abnormalities^51,52^. Clearly, long axons are a bottleneck of neuronal integrity, also because prolonged reduction of ATP can perturb transport and mitochondrial integrity in a vicious circle. We suggest that during a metabolic crisis, myelin-derived metabolites help maintaining axonal ATP to prevent irreversible damage.

## Acknowledgements

We thank A. Fahrenholz, B. Sadowski, G. Fricke-Bode and U. Kutzke for technical help, and the institute’s animal facility, the light microscopy facility, and the mechanical workshop for expert support. We also acknowledge U. Suter and the KAGS team for helpful discussions. E.A. was supported by a fellowship of the German Academic Exchange Service (DAAD). Work in the authors’ laboratory was supported by grants from the German Research Council (DFG), including SPP1757 and TRR-274. A.S.S. received support from the Cloetta Foundation and the Swiss National Science Foundation. B.P. and K.A.N. acknowledge support by the Dr. Myriam and Sheldon Adelson Medical Foundation (AMRF). K.A.N was supported by an ERC Advanced Grant (MyeliNANO).

## Author contributions

E.A., C.K., and K.A.N. conceptualized and designed the study. E.A., Z.J.L., P.D., K.K. and T.R. performed experiments. A.T., A.S.S., B.W., P.D., W.M. O.J., H.E. and J.H. supervised trainees, contributed to data analysis, and provided conceptual input. M.B., D.A., B.P. and A.B. provided mouse mutants and critical experimental advice. E.A. and K.A.N. wrote the manuscript with input from coauthors.

