## Supplementary Information for "Myelin lipids as nervous system energy reserves"

### Material and methods

#### Animals:

All mice were bred on a c57Bl6 background (except Aldh1l1-GFP) and kept under a 12h/12h day & night cycle with access to food and water *ad libitum*. Experiments were carried out in compliance with approved policies of the German state of Lower Saxony.

Transgenic mice were generated in-house by routine procedures, as previously described[1]. To visualize autophagosome in oligodendrocytes, a *mTagRFP-mWasabi-LC3* construct[2] was placed under control of the *Cnp* promoter. Genotyping was with forward (5'-CAAATAAAGCAATAGCATCACA -3') and reverse (5'- GCAGCATCCAACCAAAATCCCGG -3') primers, using the following PCR program: (30 x 58°C: 30'' ; 72°C: 45'' ; 95°C: 30'').

Eight other mouse lines were genotyped as previously published: (1) *Aldh1l1-GFP* for labeling astrocytes[3] , (2) *Cxcr-GFP* for microglia[4], (3) *Cnp-mEos2* for oligodendrocytes[1], (4) *Ng2-YFP* for OPC[5], (5) *Mfp2$^{flox/flox}$ ::Cnp-Cre*, targeting peroxisomal beta oxidation in myelinating glia[6,7], (6) *Slc2a1$^{flox/flox}$ :: Plp1$^{CreERT2}$*, targeting GLUT1 in myelinating glia[8,9], (7) *Tfeb$^{flox/flox}$ :: Cnp-Cre*, targeting autophagy in myelinating glia[7,10], and *Thy1-Ateam*, a neuronal ATP sensor[11].

The following control genotypes were used in combination with the corresponding homozygous mutants: *Cnp$^{+/+}$ :: Mfp2$^{flox/flox}$*, *Plp1$^{+/+}$ :: Slc2a1$^{flox/flox}$* + tamoxifen, and *Cnp$^{+/+}$ :: Tfeb$^{flox/ flox}$*.

#### Materials:

Reagents were purchased from Merck unless otherwise stated.

#### aCSF solution for optic nerve incubation and recording

Optic nerve incubation and electrophysiological recordings were done under constant superfusion with aCSF containing (in mM): 124 NaCl, 23 $NaHCO_3$, 3 KCl, 2 $MgSO_4$, 1.25 $NaH_2PO_4$ and 2 $CaCl_2$. aCSF was constantly bubbled with carbogen (95% $O_2$, 5% $CO_2$). A concentration of 10 mM glucose (Glc) (Sigma Aldrich, ≥99% France) was used as the standard (control). Incubation experiments were done with 0 mM glucose ("glucose-free"). For electronmicroscopy and proteomics experiments 0.5 mM and 1 mM glucose ("starvation") were applied, respectively. For electrophysiological recordings, 2.7 mM glucose was applied as a "starvation" condition unless otherwise stated. Whenever a lower glucose concentration than 10 mM was applied, the difference was substituted by sucrose (which cannot be metabolized) to maintain osmolarity.

Specific inhibitors for (i) mitochondrial beta-oxidation, 4-Bromocrotonic acid[12] (TCI, ≥98%, Japan); (ii) peroxisomal beta-oxidation, Thioridzine (Sigma Aldrich, ≥99% Germany); (iii) mitochondrial beta-oxidation of long chain fatty acids, Etomoxir[13] (Tocris, ≥98%, UK), (IV) ROS inhibitor, S3QEL-2[14] (Sigma Aldrich); (V) ROS scavenger, MitoTEMPO[15] (Sigma Aldrich) and (VI) autophagy inhibitor, Lys05[16] (Sigma Aldrich, ≥98%, China) were prepared freshly and added to aCSF at a concentration of 25, 5, 5, 10, 10 and 10 µM, respectively. To block mitochondrial oxidative phosphorylation, 5 mM sodium azide was supplemented to aCSF containing 0 mM glucose and 119 mM NaCl. The autophagy inducer, 3,4-Dimethoxychalcone[17] (DMC) (AdipoGen) was applied at 40 µM.

**Mouse optic nerve preparation and incubation**

After cervical dislocation, optic nerves were dissected before the optic chiasm and each nerve was gently removed. The prepared nerves (attached to the eyeball) were transferred into a six-well plate containing 10 ml aCSF adjusted to 37 °C. Another 90 ml aCSF was circulating during the incubation period. To study the effects of anoxia, aCSF was bubbled with nitrogen (95% $N_2$, 5% $CO_2$; Air liquide, Germany) instead of carbogen (95% $O_2$, 5% $CO_2$). To minimize the diffusion of oxygen into aCSF the wells were sealed with parafilm.

**Determining glial cell survival**

To label dying cells, optic nerves were exposed to propidium iodide (PI, 12 µM; Sigma, USA) during the last hour of incubation. Nerves were subsequently washed (10 min) in 7 ml aCSF. After 1h fixation with 4% paraformaldehyde (4% PFA in 0.1 M phosphate buffer) the nerves were detached from the eyeball and frozen blocks were prepared in Tissue-Tek® O.C.T™ compound (SAKURA, Poland). Sections (8 µm) were obtained by cryosectioning (Leica, Germany) and kept in the dark at -20 °C until further staining. Sections were washed in PBS (10 min) and stained with DAPI (1:20000 of a 1 mg/ml stock), washed again in PBS (2 x 5 min) and mounted.

All images were taken with an inverted epiflourescent microscope (Zeiss Axio Observer Z1, Germany). Illumination and exposure time settings for PI and DAPI were kept constant for all images. For the fluorescent reporter lines, different exposure times were used, according to the observed signal intensity of each fluorophore. Sections (2-3) of optic nerves were imaged and tiled arrays were stitched by the microscope software (Zen, Zeiss, Germany) for quantification.

To determine the percentage of dying cells, Fiji and Imaris softwares (version 8.1.2) were used. Stitched images were loaded in Fiji to trim areas of the optic nerves that contain dying cells unrelated to the experiment (i.e. due to normal handling). After adjusting the thresholds for each channel, single cells were automatically marked over the nucleus, manually double-checked and corrected. In the last step, co-localization of signals was calculated and data were exported (Excel files) for statistical analysis. The percentage of dying cells was obtained by dividing the number of PI over DAPI positive nuclei (PI/DAPI). For determining the frequency of each cell type, the number of fluorescent cells (GFP, YFP or mEOS2) was divided by the number of DAPI positive cells. To calculate the relative survival of each cell type, the percentage of each fluorophor-positive cell type was determined (astrocytes: Aldh1l1-GFP; microglia: Cxcr1-GFP; OPC: NG2-YFP; oligodendrocytes: Cnp-mEOS2) after the corresponding

PI/DAPI positive cells had been subtracted. The obtained values were normalized to the results for control conditions (10 mM glucose) and expressed as 'cell survival rate'. To minimize the effect of signal intensity differences between different experiment, we adjusted the threshold for quantifications based on respected controls in each experiment.

**Myelin preparation**
GLUT1 icKO mice were sacrificed five months after tamoxifen injection (age 7 months). A light weighted membrane fraction enriched in myelin was obtained from frozen half brains, using a sucrose density gradient centrifugation as previously described[18]. Briefly, after homogenizing the brains in 0.32 M sucrose solution containing protease inhibitor (cOmplete, Roche, Germany), a crude myelin fraction was obtained by density gradient centrifugation over a 0.85 M sucrose cushion. After washing and two osmotic shocks, the final myelin fraction was purified by sucrose gradient centrifugation. Myelin fractions were washed, suspended in TBS buffer (137 mM NaCl, 20 mM Tris/HCl, pH 7.4, 4°C) and supplemented with protease inhibitor (Roche, Switzerland).

**Western blotting**
Immunoblotting and fast green staining was performed as previously described[19] using the following primary and secondary antibodies: ACAT1 (1:3000, 16215-1-AP, protein tech), BDH1 (1:500, 15417-1-AP, proteintech), LC3B (1:2000, NB100-2220, Novusbio), Na+/K+ATPaseα1 (1:1000, Abcam), GLUT1 (1:1000)[20], GLUT2 (1:1000, ab54460, abcam), GLUT3 (1:1000, ab191071, abcam), GLUT4 (1:1000, 07-1404, Millipore), MCT1 (1:1000)[21], carbonic anhydrase 2 (CA2, 1:1000)[22], and α-tubulin (TUBA, 1:1000, T 5168, Sigma, 1:10000, Mouse IgG H&L Antibody Dylight™ 680 Conjugated, 610-144-002; Rabbit IgG H&L Antibody DyLight™ 800 Conjugated, 611-145-002; Rockland). Signal intensities, analyzed with the Image Studio software Licor or Fiji, were normalized to the corresponding total protein load, which were quantified by fast green staining. Obtained values were normalized to the mean of the respective values from control mice.

**Proteome analysis and Western blots**
Optic nerves were collected after incubation in aCSF with 10 mM glucose (control), 0 mM or 1 mM glucose, transferred to microtubes, and kept at -80 °C until further analysis. To minimize variability, one nerve from a mouse was incubated under starvation or glucose-depreviation conditions, and the other under the control condition.  Nerves from two mice were pooled for protein extraction, homogenized in 70 µl of RIPA buffer (Tris-HCl 50 mM; Sigma, USA), Na-deoxycholate (0.5%; Sigma Aldrich, New Zealand), NaCl (150 mM), SDS (0.1%; Serva, Germany), Triton X100 (1%; Sigma, USA), EDTA (1 mM) and complete protease inhibitor cocktail (Roche, Switzerland) by using ceramic beads in a Precellys homogenizer (for 3 x 10sec at 6500rpm) (Precellys 24, Bertin Instruments, USA). After a 5min centrifugation at 13000 rpm at 4 °C, the supernatant was used for protein determination (DC Protein Assay reagents, Bio Rad, USA) according to the manufacturer protocol, with the absorbance of samples at 736 nm

(Eon microplate spectrophotometer, Biotek Instruments, USA). Proteins (0.5 µg) were separated on 12% SDS-PAGE gels and subjected to silver staining [23].

**Proteomics**

Proteome analysis of purified myelin was performed as recently described[18,19] and adapted to optic nerve lysates[24]. Briefly, supernatant fractions corresponding to 10 µg protein were dissolved in lysis buffer (1% ASB-14, 7 M urea, 2 M thiourea, 10 mM DTT, 0.1 M Tris pH 8.5). After removal of the detergents and protein alkylation proteins were digested overnight at 37°C with 400 ng trypsin. Tryptic peptides were directly subjected to LC-MS-analysis. For quantification according to the TOP3 approach[25], aliquots were spiked with 10 fmol/µl of Hi3 EColi standard (Waters Corporation, USA), containing a set of quantified synthetic peptides derived from E. coli. Peptide separation by nanoscale reversed-phase UPLC was performed on a nanoAcquity system (Waters Corporation, USA) as described[24]. Mass spectrometric analysis on a quadrupole time-of-flight mass spectrometer with ion mobility option (Synapt G2-S, Waters Corporation) was performed in UDMSE[26] and MSE mode as established for proteome analysis of purified myelin[19,27] to ensure correct quantification of myelin proteins that are of high abundance. Processing of LC-MS data and searching against UniProtKB/Swiss-Prot mouse proteome database was performed using the Waters ProteinLynx Global Server (PLGS) version 3.0.3 with published settings[19]. For post-identification analysis including TOP3 quantification of proteins, the freely available software ISOQuant[26] (www.isoquant.net) was used. False discovery rate for both peptides and proteins was set to 1% threshold and only proteins reported by at least two peptides (one of which unique) were quantified as parts per million (ppm) abundance values (i.e. the relative amount (w/w) of each protein in respect to the sum over all detected proteins). The Bioconductor R packages "limma" and "q-value" were used to detect significant changes in protein abundance by moderated t-statistics as described[28]. Optic nerve fractions from five animals per condition (10 mM glucose vs. 1 mM glucose/9 mM sucrose; 10 mM glucose vs. 0 mM glucose/10 mM sucrose) were processed with replicate digestion, resulting in two technical replicates per biological replicate and thus in a total of 20 LC-MS runs to be compared per individual experiment.

**Electron microscopy**

Freshly prepared or incubated optic nerves were immersion fixed in 4 % formaldehyde, 2.5 % glutaraldehyde (EM-grade, Science Services, Munich, Germany) and 0.5 % NaCl in phosphate buffer pH 7.4 overnight at 4 °C. Fixed samples were embedded in EPON after dehydration with acetone as previously described[29]. Sections of 50-60 nm thickness were obtained with the Leica UC7 ultramicrotome (Leica, Vienna, Austria) equipped with a diamond knife (Histo 45° and Ultra 35 °C, Diatome, Biel, Switzerland) and imaged using a LEO EM 912AB electron microscope (Zeiss, Oberkochen, Germany) equipped with an on-axis 2048x2048-CCD-camera (TRS, Moorenweis, Germany).

**EM analysis**

EM images from optic nerves were imported into the Fiji software. In order to create an unbiased selection of axons for which the g-ratio was calculated, a grid consisting of 1 $\mu m^2$ squares for ex vivo and 4 $\mu m^2$ squares for in vivo experiment was overlaid on each image and axons that were crossed by the intersecting lines were selected for quantification, with the requisites of 1) the axon being in focus and 2) the axon shape being not evidently deformed. For each axon, three circles were manually drawn around the axonal membrane, the inner layer, and the outer layer of myelin. When myelin was not evenly preserved, the myelin thickness of the adjacent, preserved area was used as a proxy and the circle was corrected accordingly. The obtained area (A) of each circle was converted into the corresponding diameter using the formula $A = \pi r2$ and the g-ratio (outer diameter/axon diameter) was calculated. In ex vivo experiments, the obtained data from the axons with a diameter smaller than 2 µm were used for further analysis. The mean g-ratio for axons from one nerve was used for statistical analysis and presented as a datapoint in the bar graphs.

To determine the axonal size distribution in the optic nerves, axons were binned by increasing caliber and the number of counted axons for each caliber bin was divided by the total numbers of counted axons per nerve and presented as a single data point.

In optic nerves, the total number of axons, the number of degenerating and unmyelinated axons was counted in microscopic subfields of 1445 square µm. Using these data, the percentage of degenerating and unmyelinated axons was calculated.

For nerves incubated in vitro, the percentage of axons containing vesicle-like structures in glial cytoplasm underneath their myelin sheath the frequency was also determind.

**Electrophysiological recording**

All mice used for optic nerve electrophysiology were 8-12 weeks old (unless otherwise stated). Recordings were performed as described previously[11,30,31]. Briefly, optic nerves were carefully dissected and quickly transferred into the recording chamber (Harvard Apparatus, Holliston, USA) and continuously superperfused with aCSF. A temperature controller (TC-10, NPI electronic, Tamm, Germany) maintained the temperature at 37 °C.

To assure the optimal stimulation and recording condition, custom-made suction electrodes were back-filled with aCSF containing 10 mM glucose. To achieve supramaximal stimulation, a battery (Stimulus Isolator 385; WPI, Berlin, Germany) was used to apply a current of 0.75 mA at the proximal end of the optic nerve to evoke a CAP at the distal end acquired by the recording electrode at 100 kHz connected to an EPC9 amplifier (Heka Elektronik, Lambrecht/Pfalz, Germany). The signal was pre-amplified 10 times using Ext 10-2F amplifier (NPI electronic) and further amplified (20-50 times) and filtered at 30kHz, using a low-noise voltage preamplifier SR560 (Stanford Research System). All recordings were done after nerve equilibration for 2h (except for excitability and nerve conduction velocity measurments) in aCSF containing 10 mM glucose, during which the CAP was monitored every 30 sec until a stable waveform was reached.

To measure the excitability of nerves, CAPs were evoked with currents starting from 0.05 mA and (using 0.05 mA steps) increased to 0.75 mA. The analysed CAP area (CAPA) for each current was normalized to the obtained CAP at 0.75 mA. Nerve conduction velocities (NCV) of optic nerves were determined by dividing the length of each nerve to the latency of second peak for each nerve.

**Confocal imaging acquisition**

An up-right confocal laser scanning microscope (Zeiss LSM 510 META/NLO, Zeiss, Oberkochen, Germany) equipped with an Argon laser and a 63x objective (Zeiss 63x IR-Achroplan 0.9 W) was used for live imaging of the optic nerve for ATP measurement as reported previously[11,32]. The immersion objective was placed into superfusing aCSF on top of the clamped optic nerve with electrodes and images acquired with the time resolution of 30 sec. A frame size of 114.21 x 133.30 µm (pinhole opening: 168 µm, pixel dwell time: 3.66 µs) was scanned (2x averaging) for CFP (Ex 458 nm; Em 470–500 nm), FRET (Ex 458 nm; Em long pass 530 nm) and YFP (Ex 514 nm; Em long pass 530 nm) channel and the focus was adjusted manually based on eye estimation of the nerve movement. To image autophagosome formation in the mouse optic nerve (2-5 month-old), suction electrodes were used for fixing the nerve. A frame size of 133.45 x 76.19 µm was scanned for RFP (Ex 543 nm; Em 565-615 nm) and Wasabi (Ex 488 nm ; Em BP 500-550 nm) channels with pinhole adjusted at 384 µm and pixel dwell time: 58.4 µs.

**CAP analysis**

Optic nerve function can be measured quantitatively by calculating the area underneath the evoked waveform. This `compound action potential area' (CAPA) represents the conduction of nearly all optic nerve axons. The evoked waveform from the optic nerve includes three peaks that are representative of different axons with different rate of signal speed[30,31].

The time between the first peak of the CAP waveform (at approx. 1.2 ms following the stimulation, depending on the electrodes used) and the end of third peak (depending on the nerve length) at the last few minutes of the baseline recording was defined as the time range for CAPA integration. This window was kept constant for all recorded traces for each nerve. The calculated CAPA was then normalized to the average obtained from the last 30 min of baseline recordings. The results from several nerves were pooled, averaged and after binning plotted against time. Bar graphs depict the average CAPA for short time windows or the calculated CAPA for larger time windows. Overall nerve conductivity was determined from the 'CAPA area', i.e. the area under multiple CAPA curves obtained for the nerves in the experimental arm after normalizing these readings to the corresponding mean values from control nerves.

**ATP quantification**

The relative level of ATP was calculated as previously reported[11]. Images were loaded in Fiji and the area of the nerve that was stable during the imaging was selected for measuring the mean intensity for three different channels: FRET, CFP and YFP. The FRET /CFP ratio was calculated to give a relative ATP concentration. The ratio was normalized to 0 and 1.0 by using the values obtained during the phase of mitochondrial blockade/ glucose deprivation (MB+GD, 5 mM Azide, 0 mM glucose) and baseline (10 mM glucose) respectively. Bar graphs were obtained by averaging the values obtained at the last 5min of each step of applied protocol for electrophysiology recordings of each nerve.

**Immunohistology**

Longitudinal cryosections of the optic nerve were fixed in 4% PFA (10-20 min) followed by washing in PBS (3x5 min). After 30 min permeabilization with 0.4% Triton in PBS at room temperature (RT), blocking was performed for 30 min at RT in blocking solution (4% horse serum, 0.2% triton in PBS). Incubation with primary antibody (Iba1, 1:1000, Wako) was performed in blocking solution (1% HS, 0.02% triton in PBS) at 4 °C. After washing with PBS (3x10 min), sections were incubated with secondry antibody in PBS/BSA containing DAPI (1:2000, stock 1 mg/ml) for 1h at RT. The washed sections (3x10 min) PBS were mounted and microscopy was performed.

**Autophagosome quantification**

Microscopy images were processed with Fiji software and the number of autophagosomes (appearing as puncta) and the number of cell bodies in each image were manually quantified. The total numbers of counted autophagosomes were divided by the total numbers of cell bodies in the same images. At least three images from different regions of the nerve were analyzed for each nerve and the average of the obtained values was presented as a single data point.

**Mouse behavior**

All measurments were performed by the same experimentator, blinded to the animals' genotype. Mice were trained weekly for six weeks before actual testing. For the Rotarod test, mice were placed on the horizontal rod. Rotation started with 1 rpm and one unit (rpm) was added every 10 seconds. The rpm value at which mice fell was recorded and the average of three repeats was reported for each mouse.

For grip strength measurements, mice were allowed to grasp a metal bar that connected to the grip strength meter. Holding on with their forelimbs, mice were slowly pulled backward until the grip was lost. The average of three measurments was reported as a data point.

**Data presentation and statistics**

All data are presented as mean ± SEM. For cell death measurements in the optic nerve, the quantification of 2-3 sections (except for 16h incubation (Fig.1c) that was 1-3 sections) from the same nerve were combined and the mean was taken as one data point. The n-number indicates the total number of optic nerves analyzed for each condition. For 1 mM glucose experiments two nerves were pooled for the analysis. The data were analyzed using t-test (unpaired, two tailed distribution, two-sample equal variance/homoscedastic).

### Supplementary Images

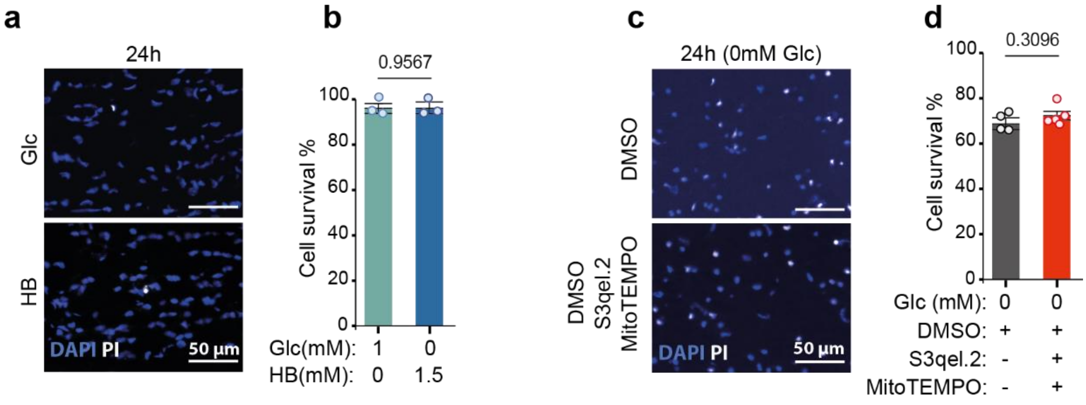


**Suppl Fig 1 Death of glucose-deprived optic nerve glial cells is not caused by oxidative stress**

**a,** Acutely isolated mouse optic nerves (age 2 months) were maintained for 24h either in the presence of low (1mM) glucose or 1.5mM beta-hydroxybutyrate (HB). Longitudinal nerve sections were stained with DAPI (blue) and propidium iodide (in white).

**b,** Bar graphs showing the percentage of viable cells for each condition (compare to Fig 1c). Normal survival is possible in the presence of 1mM glucose or 1.5mM HB, indicating the critical role of glucose in energy production (n = 3 each; mean+/-SEM, t-test).

**c,** Longitudinal sections of optic nerves stained with PI and DAPI after 24h incubation in glucose-free aCSF and in the presence of 10 µM of S3qel-2 (inhibitor for ROS production in complex III of electrone transport chain) and 10 µM MitoTEMPO (mitochondrial ROS scavenger) or vehicle (DMSO) as control.

**d,** Quantified cell survival (PI/DAPI) indicates that ROS do not contribute to cell death (n=4 for control and n=5 for ROS inhibitor/scavenger; wild type mice, 8-10 weeks old; t-test, mean +/-SEM).

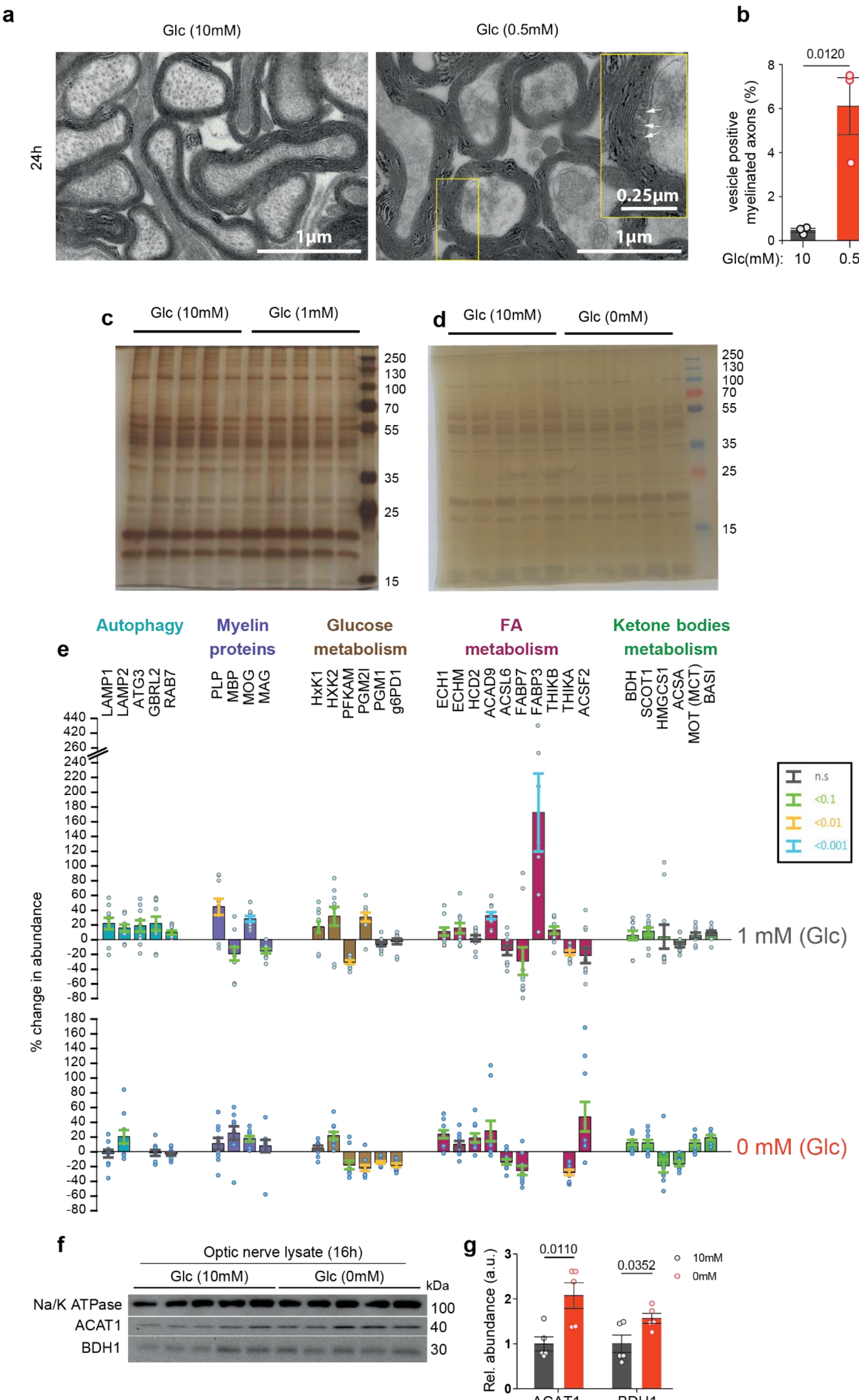

a
Glc (10mM)
Glc (0.5mM)
24h
1µm
0.25µm
b
0.0120
vesicle positive myelinated axons (%)
Glc(mM): 10 0.5
c
Glc (10mM)
Glc (1mM)
250
130
100
70
55
35
25
15
d
Glc (10mM)
Glc (0mM)
e
Autophagy
Myelin proteins
Glucose metabolism
FA metabolism
Ketone bodies metabolism
LAMP1 LAMP2 ATG3 GBRL2 RAB7
PLP MBP MOG MAG
HxK1 HXK2 PFKAM PGM2I PGM1 g6PD1
ECH1 ECHM HCD2 ACAD9 ACSL6 FABP7 FABP3 THIKB THIKA ACSF2
BDH SCOT1 HMGCS1 ACSA MOT (MCT) BASI
% change in abundance
n.s
<0.1
<0.01
<0.001
1 mM (Glc)
0 mM (Glc)
f
Optic nerve lysate (16h)
Glc (10mM)
Glc (0mM)
kDa
Na/K ATPase
ACAT1
BDH1
100
40
30
g
Rel. abundance (a.u.)
0.0110
0.0352
10mM
0mM
ACAT1
BDH1

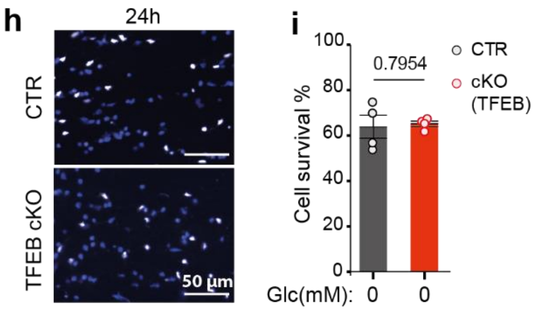


**Suppl Fig 2 Upregulation of autophagy and lipid metabolism in metabolically stressed optic nerves**

**a,** Electron micrographs of optice nerves, incubated for 24 in 10mM glucose (left) or 0.5mM glucose (right). Starved nerves reveal periaxonal vesicular structures (white arrows) indicating myelin degradation under low glucose condition (age 2 months, wild type).

**b,** Quantification of the data in (a) comparing the percentage of myelinated axons with periaxonal vesicular structures in cross section (n=3 for control and n=3 for low glucose condition; t-test, mean +/-SEM).

**c,** Silver stained protein gel of optic nerve lysates, prepared after 24h incubation in medium with 10 mM glucose (left) or 1 mM glucose (right). Each lane is a pool of 2 wildtype nerves (age 2 months).

**d,** Silver stained protein gel, prepared after 16h incubation in 10 mM glucose (left) or 0 mM glucose (right).

**e,** Relative abundance of selected optic nerve proteins in lysates after 24h in medium with 1 mM glucose (top row) or 16h in 0 mM glucose (bottom row). Quantification is in comparison to nerves maintained in 10 mM glucose for 24 or 16 h. Enzymes of glucose and lipid metabolism show only moderate changes in abundance. Autophagy related proteins are only increased in the presence of 1mM glucose, indicating the requirement of glucose for RNA synthesis and protein expression. (n=5, two technical replicates each; moderated t-statistics; Mean +/-SEM).

**f,** Western blots of lysates from wildtype optic nerves incubated *ex vivo* in 10mM or 0mM glucose for 16h (age 8-12 weeks old, n=5 for each condition).

**g,** Quantification of the data in (f) for ACAT1 and BDH1, normalized to protein input (t-test; mean +/-SEM).

**h,** Cell survival analysis of 24h glucose-deprived optic nerves from TFEB cKO mice. Images from longitudinal sections were stained with PI and DAPI.

**i,** Quantified data from (h). There is no difference of cell survival in optic nerves from wildtype (n=4) and TFEB cKO mice (n=4) (age 8-12 weeks old; t-test, mean+/-SEM).

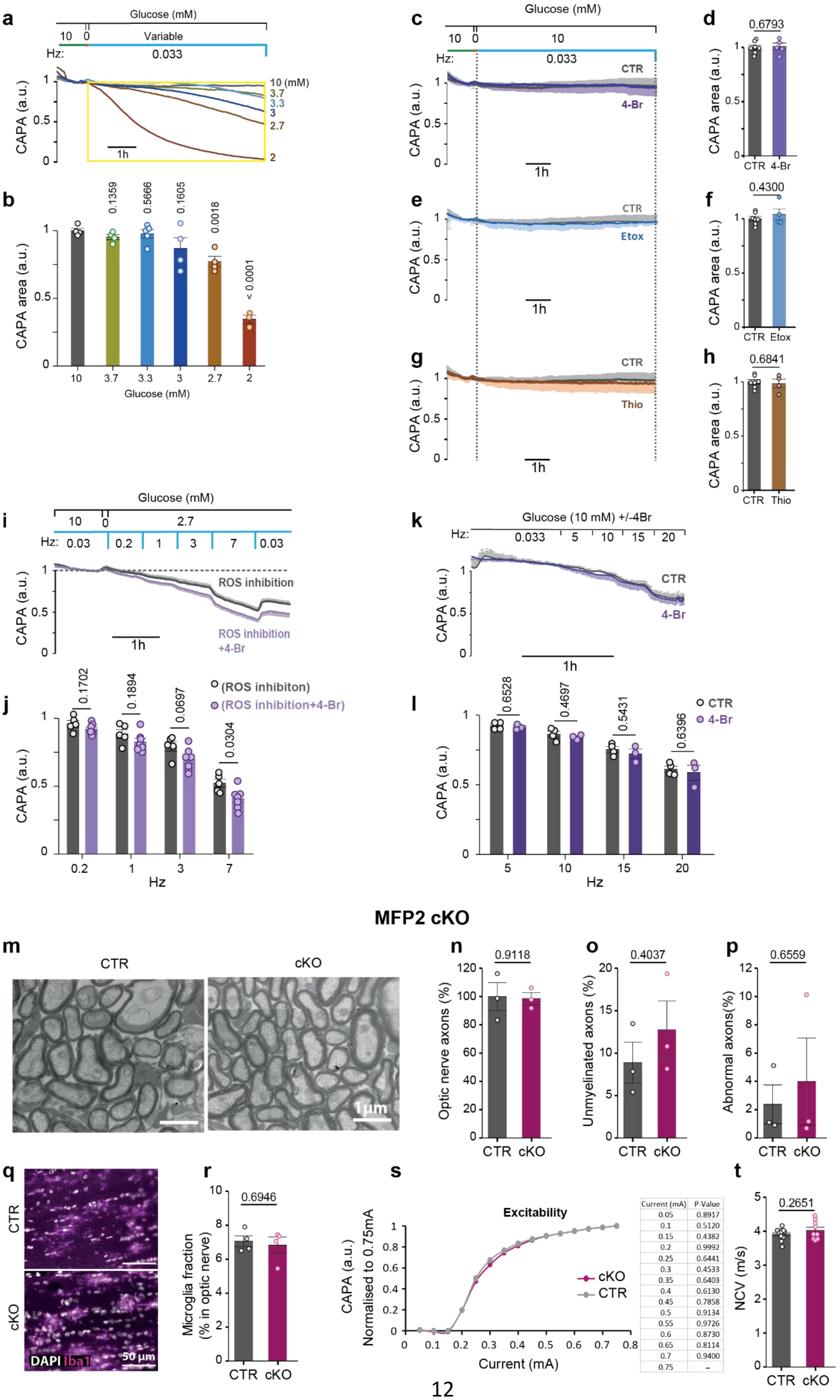


| Current (mA) | P-Value |
|---|---|
| 0.05 | 0.8917 |
| 0.1 | 0.5120 |
| 0.15 | 0.4382 |
| 0.2 | 0.9992 |
| 0.25 | 0.6441 |
| 0.3 | 0.4533 |
| 0.35 | 0.6403 |
| 0.4 | 0.6130 |
| 0.45 | 0.7858 |
| 0.5 | 0.9134 |
| 0.55 | 0.9726 |
| 0.6 | 0.8730 |
| 0.65 | 0.8114 |
| 0.7 | 0.9400 |
| 0.75 | - |

**Suppl Fig 3 Fatty acid beta-oxidation in oligodendrocytes supports conduction of starved optic nerves**

**a,** Empirical determination of a minimal glucose concentration in artificial cerebrospinal fluid (aCSF) at which optic nerves can maintain a low spiking frequency (1/30s) for at least 3 hours. Compound action potentials (CAP) were recorded and areas underneath the curve (CAPA) normalized to control values obtained with 10mM glucose.

**b**, Bar graphs of the CAPA summation (CAPA area) calculated for the time window defined by yellow box for different glucose concentrations using the same data as in (a) after normalization to control condition. (Error bars: mean+/-SEM, t-test).

**c,** Optic nerve recordings in aCSF+10mM glucose +/-25 µM 4-Br (n=7 for control and n=5 for 4-Br treated nerves; wild type mice, 8-12 weeks old).

**d,** Bar graph of the CAPA summation (CAPA area) for the defined time window (dash lines) in (c). values were normalized to control (error bars: mean+/-SEM, t-test).

**e,** Normal conduction of optic nerves kept in aCSF with 10mM glucose and 5 µM Etomoxir (Etox; n=4) in comparison to 10mM glucose control (same data as in c). Wildtype mice, age 8-12 weeks.

**f,** Bar graph comparing the calculated CAPA summation (CAPA area) for the time window between the dash lines in (e). obtained values were normalized to control condition (mean+/-SEM, t-test).

**g,** Normal conduction of optic nerves kept in aCSF with 10mM glucose and 5 µM Thioridazine (Thio; n=7 in comparison to 10mM glucose control (same data as in c). Wildtype mice, age 8-12 weeks.

**h,** Bar graph comparing the calculated CAPA summation (CAPA area) for the depicted time window (dash lines) in (g). obtained values were normalized to control (mean+/-SEM, t-test).

**i,** Conductivity of optic nerves under low glucose (aCSF+2.7mM glucose +/-4-Br(25µM)) in the presence of the ROS production inhibitor (S3qel-2, 10µM) and mitochondrial ROS scavenger (MitoTEMPO, 10µM). n=5 for control (without 4-Br) and n=6 (plus 4-Br); wild type mice, 8-12 weeks old.

**j,** Bar graph comparing calculated CAPA from the data in (i) for the average of CAPA recorded during the last 5 min of each step of the RAMP protocol. Mean+/-SEM, t-test.

**k,** Optic nerves spiking at high frequencies. Recordings were in aSCF (10mM glucose) +/- 25 µM 4-Br, using a RAMP protocol of increasing frequencies between 5 and 20Hz (n=4 for control and n=3 for 4-Br treated nerves). Wild type mice, age 8-12 weeks.

**l,** Bar graphs of CAPA, calculated for each nerve over 5 min at the indicated frequencies, using recordings in (k). Mean+/-SEM, t-test.

**m,** Electrone microscopic images from optic nerve cross sections of MFP2 cKO mice.

**n,** Bar graphs comparing the number of axons (in m) in MFP2 cKO mice normalized to control nerves (n=3 each). Age 8-12 weeks; mean+/-SEM, t-test.

**o,** Bar graphs comparing the number of unmyelinated axons (in m) in MFP2 cKO mice normalized to control nerves (n=3 each). Age 8-12 weeks; mean+/-SEM, t-test.

**p,** Percentage of axons with abnormal morphology (same images in m) in MFP2 cKO mice normalized to control nerves (n=3 each). Age 8-12 weeks; mean+/-SEM, t-test.

**q,** longitudinal optic nerve section from MFP2 cKO mice and controls immunostained for Iba1 and counterstained with DAPI.

**r,** Bar graph quantifying Iba+ microglia in the optic nerve, normalized to DAPI+ nuclei (from images in q) n=4 for control and n=5 for MFP2 cKO. Age 8-12 weeks (mean+/-SEM, t-test).

**s,** Normal excitability of optic nerves from MFP2 cKO mice (n=11 for CTR and n=7 for cKO; t-test).

**t,** Bar graph comparing optic nerve conduction velocity (NCV) determined by deviding the latency of the second CAP peak after stimulation to the length of nerve (n=11 for control and n=9 for cKO; 8-12 weeks old; mean+/-SEM, t-test; common data with s).

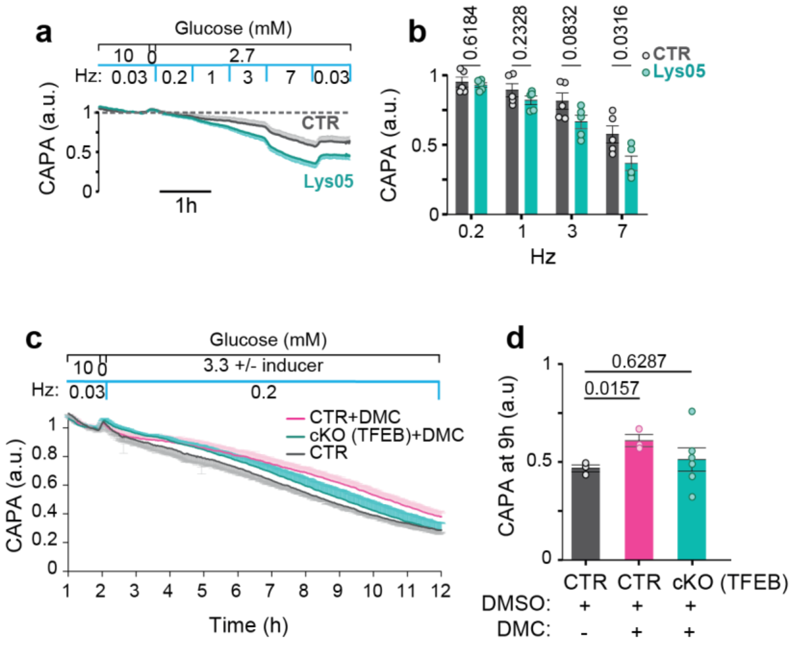


**Suppl Fig 4 Preexisting autophagy contributes to oligodendroglial support of axonal conduction in starved optic nerves**

**a,** CAP recordings from wildtype optic nerves kept under low glucose condition (aCSF with 2.7mM glucose) in the presence or absence of 10 µM autophagy inhibitor Lys05 (8-12 weeks old wild type mice; n=5 each).

**b,** Bar graph representing the average CAPA during the last 5 min of each stimulation step (same data in a; mean+/-SEM, t-test).

**c,** Effect of the autophagy inducer DMC (40 µM) on nerve conduction under low glucose condition. Note that the inducer improves CAPA in wildtype optic nerves but in nerves from TFEB cKO mice.

**d,** Quantification of the data in (c) with a comparison of CAPA at 9h (average of 5min recordings). Statistics: n=3 for control (+DMSO), n=3 for control +DMC, and n=6 for TFEB+DMC. 3-5 months old mice (mean+/-SEM, t-test).

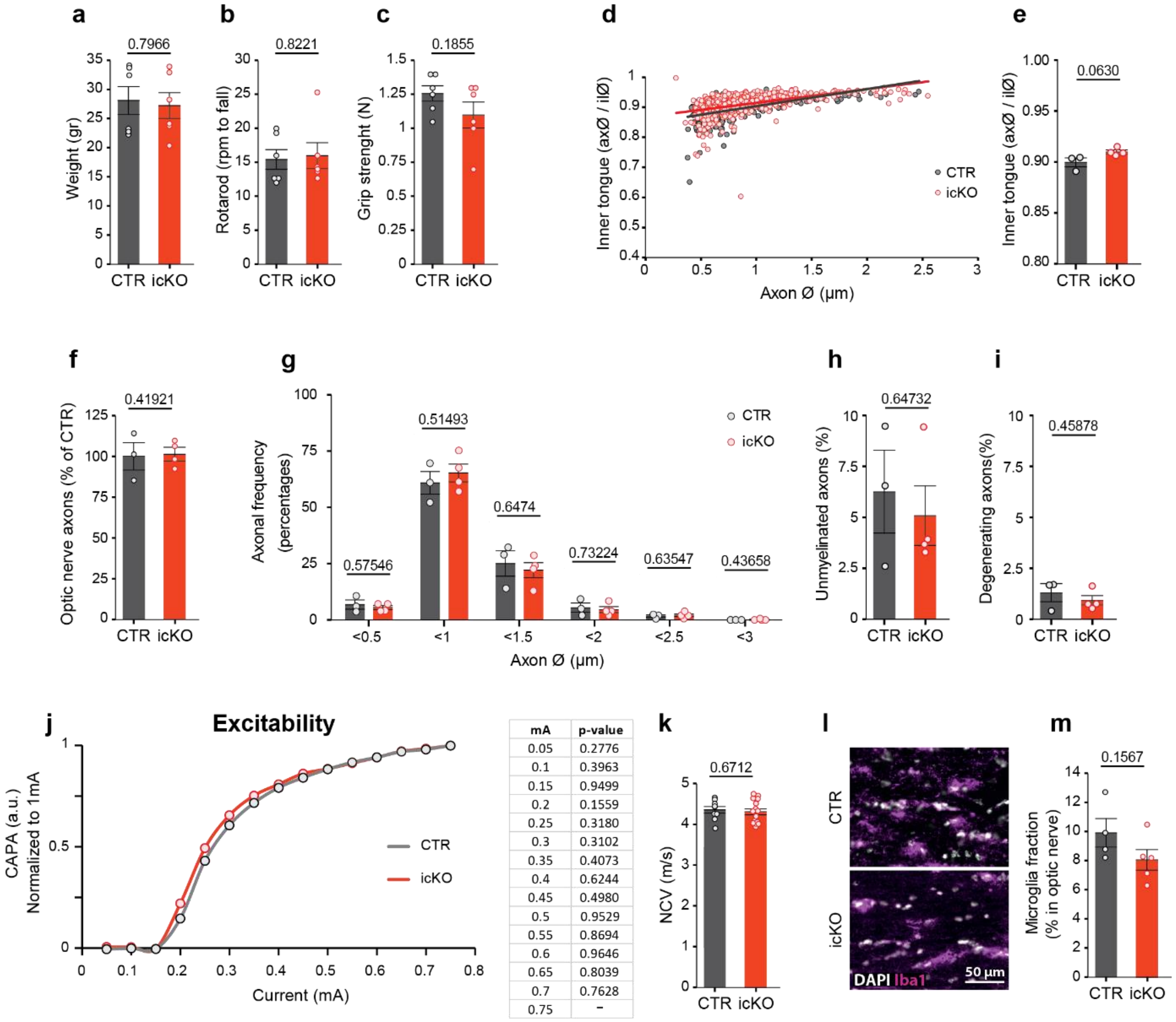


| mA | p-value |
|---|---|
| 0.05 | 0.2776 |
| 0.1 | 0.3963 |
| 0.15 | 0.9499 |
| 0.2 | 0.1559 |
| 0.25 | 0.3180 |
| 0.3 | 0.3102 |
| 0.35 | 0.4073 |
| 0.4 | 0.6244 |
| 0.45 | 0.4980 |
| 0.5 | 0.9529 |
| 0.55 | 0.8694 |
| 0.6 | 0.9646 |
| 0.65 | 0.8039 |
| 0.7 | 0.7628 |
| 0.75 | - |

**Suppl Fig 5 Hypomyelinated optic nerves in normally behaving GLUT1 icKO mice lack histological signs of pathology or gliosis.**

**a,** Normal body weight of GLUT1 icKO mice at the age of 7 months (5 months post tamoxifen). (n=6 for control and n=6 for icKO; mean+/-SEM, t-test).

**b,** Normal rotarod performance of GLUT1 icKO mice shown as the speed (rpm) at which the mice fall (same mice in a; mean+/-SEM, t-test).

**c,** Normal forelimb grip strength of GLUT1 icKO mice (same mice as in a; mean+/-SEM, t-test).

**d,** Quantification of the inner tongue size as a function of axon caliber, by plotting the axon diameter (ax) by the respective diameter of a circle defined by the inner surface of the compacted myelin sheath (il), analogous to g-ratios (same images in fig4 f).

**e,** Tendency for smaller average inner tongue sizes in GLUT1 icKO mice. (same data as in d; n = 3 for control and n=4 for icKO mice; mean+/-SEM, t-test).

**f,** Normal number (density) of optic nerve axons in GLUT1 icKo mice normalized to controls (same images from fig4 f; mean+/-SEM, t-test).

**g,** Normal axon size distribution in optic nerve axons from GLUT1 icKo mice (same images from fig4 f; mean+/-SEM, t-test).

**h,** Normal percentage of unmyelinated axons in GLUT1 icKO mice (same images from fig4 f; mean+/-SEM, t-test).

**i,** Normal percentage of axons showing morphological abnormalities by EM analysis (from images in fig4 f; mean+/-SEM, t-test).

**j,** Normal excitability of optic nerves from GLUT1 icKO mice 4-5 mongths after recombination, recorded with increasing current of stimulation. Calculated CAPA for each current was normalized to recorded CAPA at 0.75mA (n=6 for control and n=12 for icKO; t-test).

**k,** Normal optic nerve conduction velocity (NCV) of GLUT1 icKO mice, calculated by deviding latency of the second CAP peak to the length of the nerve (common data with j; n=9 for control and n=15 for icKO; mean+/-SEM, t-test).

**l,** Longitudinal sections of GLUT1 icKO optic nerves immunostaned for Iba1 and counterstained with DAPI (4-5 months post tamoxifen).

**m,** Normal number of Iba1+ microglia in optic nerve from GLUT1 icKO mice, normalized to the number of DAPI+ nuclei in the same area (same images in l). Controls n=4, GLUT1 icKO n=5 (mean+/-SEM, t-test).
